# Sodium nitroprusside prevents the detrimental effects of glucose on the neurovascular unit and behaviour in zebrafish

**DOI:** 10.1101/576942

**Authors:** K. Chhabria, A. Vouros, C. Gray, R.B. MacDonald, Z. Jiang, R.N. Wilkinson, K Plant, E. Vasilaki, C. Howarth, T.J.A. Chico

**Affiliations:** Neuroimaging in Cardiovascular Disease (NICAD) network, University of Sheffield; Department of Infection, Immunity and Cardiovascular Disease, University of Sheffield Medical School, Beech Hill Road, Sheffield S10 2RX United Kingdom.; The Bateson Centre, Firth Court, University of Sheffield, Western Bank, Sheffield, S10 2TN United Kingdom.; Department of Psychology, University of Sheffield, Cathedral Court, 1 Vicar Lane, Sheffield, S1 2LT United Kingdom.; Department of Computer Science, University of Sheffield, Portobello, S1 4DP United Kingdom.

**Keywords:** Neurovascular unit, behaviour, nitric oxide, hyperglycemia, zebrafish

## Abstract

Diabetes is associated with dysfunction of the neurovascular unit, although the mechanisms of this are incompletely understood, and currently no treatment exists to prevent these negative effects. We previously found that the NO donor sodium nitroprusside (SNP) prevents the detrimental effect of glucose on neurovascular coupling in zebrafish. We therefore sought to establish the wider effects of glucose exposure on both the neurovascular unit and on behaviour in zebrafish and the ability of SNP to prevent these.

We incubated 4 days post fertilisation (dpf) zebrafish embryos in 20mM glucose or mannitol for five days until 9dpf, with or without 0.1mM SNP co-treatment for 24h (8-9dpf), and quantified vascular nitric oxide reactivity, vascular mural cell number, expression of a *klf2a* reporter, glial fibrillary acidic protein (GFAP) and TRPV4, as well as spontaneous neuronal activation at 9dpf, all in the optic tectum. We also assessed the effect on light/dark preference and locomotory characteristics during free-swimming studies.

We find that glucose exposure significantly reduced nitric oxide reactivity, *klf2a* reporter expression, vascular mural cell number and TRPV4 expression, while significantly increasing spontaneous neuronal activation and GFAP expression (all in the optic tectum). Furthermore, when we examined larval behaviour we found glucose exposure significantly altered light/dark preference and high and low speed locomotion while in light. Co-treatment with SNP reversed all these molecular and behavioural effects of glucose exposure.

Our findings comprehensively describe the negative effects of glucose exposure on the vascular anatomy, molecular phenotype, and function of the optic tectum and on whole organism behaviour. We also show that SNP or other NO donors may represent a therapeutic strategy to ameliorate the complications of diabetes on the neurovascular unit.

## Introduction

The prevalence of diabetes has quadrupled in the previous two decades, incurring an enormous burden of morbidity and healthcare expenditure worldwide (Zhang *et al*., 2010; Scully, 2012). Diabetes is a risk factor for both macrovascular (such as myocardial infarction and stroke) and microvascular (causing renal impairment and retinopathy) disease (Chase *et al*., 1989; Jorgensen *et al*., 1994; Miettinen *et al*., 1998; Stratton *et al*., 2000). Diabetes is also associated with neurological disorders including cognitive impairment (dementia) (Stewart & Liolitsa, 1999; Areosa & Grimley, 2002; MacKnight *et al*., 2002; Ciudin *et al*., 2017; Simo *et al*., 2017; Groeneveld *et al*., 2018). The mechanisms underlying this association are incompletely understood, and no specific therapies have been identified to prevent or reverse the effects of diabetes on neurological function.

Both diabetes and neurological diseases are associated with dysfunction of the neurovascular unit (NVU) (Zlokovic, 2010; Mogi & Horiuchi, 2011; Gardner & Davila, 2017). The NVU comprises neurons, astrocytes, myocytes, pericytes, endothelial cells (ECs) and extracellular matrix. Interactions between these ensures neuronal energy demands are met through increased local blood flow via neurovascular coupling (NVC) (Roy & Sherrington, 1890; Attwell *et al*., 2010). Recent evidence suggests that ECs are crucial to NVU function (Toth *et al*., 2015; Guerra *et al*., 2018) as they release vasoactive substances such as nitric oxide (NO) (Ignarro *et al*., 1987; Palmer *et al*., 1987). NO production is regulated by various endothelial genes including the Kruppel-like family of transcription factors (KLFs)(Gracia-Sancho *et al*., 2011) particularly KLF2 which is regulated by changes in flow and inflammation (Dekker *et al*., 2002; SenBanerjee *et al*., 2004b). ECs share a common basement membrane with pericytes that aid EC development (Gerhardt & Betsholtz, 2003). Pericyte coverage, essential to both blood brain barrier integrity and NVC, is affected in various neuropathologies (Tilton *et al*., 1985; Frank *et al*., 1990; Peppiatt *et al*., 2006; Pfister *et al*., 2008; Vates *et al*., 2010; Sagare *et al*., 2013; Hall *et al*., 2014). Astrocytes are the predominant glial cell in the brain and perform several functions including release of vasoactive factors (Zonta *et al*., 2003; Takano *et al*., 2006). Astrocytes also express glutamine synthetase (GS), an enzyme involved in the recycling of glutamate released by active neurons (Bergles *et al*., 1999; Bringmann *et al*., 2013). Mammalian studies show that astrocytes sense changes in vascular tone through activation of the mechanosensor TRPV4 (Vallinoid transient receptor potential) (Filosa *et al*., 2013), which is also expressed in ECs. Together all these cell types maintain the functional NVU.

The zebrafish is increasingly used as a model of human disease (Dooley & Zon, 2000; Lieschke & Currie, 2007). This has a number of advantages over existing mammalian models, particularly ease of *in vivo* cellular imaging and the ability to test the effect of drugs by immersion. Although most zebrafish studies attempting to model human disease have examined the anatomic or molecular effects of genetic or other manipulation (Patton & Zon, 2001; Lieschke & Currie, 2007), a range of behavioural assays have been applied to study more complex “whole organism” phenotypes, such as memory or aggression (Blaser & Gerlai, 2006; Oliveira *et al*., 2011). The zebrafish has previously been used as a model of diabetes, either by exposure to medium containing glucose (Capiotti *et al*., 2014), genetic manipulation (Kimmel *et al*., 2015) or ablation of the beta-cells of the pancreas (Pisharath *et al*., 2007).

We recently established a novel zebrafish larval model of NVC in which incubation of larvae in 20mM glucose impaired NVC. We found the NO donor sodium nitroprusside (SNP) rescued this effect (Chhabria *et al*., 2018) although the mechanism for this is unclear. We therefore wish to better understand the mechanism and consequences of NVU dysfunction induced by glucose. In the present study we have now examined the effect of glucose exposure on NO production in the NVU, *klf2a* expression, mural cell number, glial fibrillary acidic protein (GFAP) expression, TRPV4 expression, spontaneous neuronal activation, light/dark preference and larval locomotory behaviour. We find that glucose exposure affects all these aspects of NVU function and behaviour, and that co-treatment with SNP completely prevents all the detrimental effects of glucose exposure. Our findings provide insight into the effect of hyperglycaemia on NVU function and further support for the possibility that NO donors represent plausible drug candidates to ameliorate the detrimental effects of hyperglycemia.

## Materials and Methods

### Transgenic zebrafish

All zebrafish studies were conducted in accordance with the Animals (Scientific Procedures) Act, 1986, United Kingdom and covered by Home Office Project Licence 70/8588 held by TC. Reporting of experimental outcomes were in compliance with ARRIVE (Animal Research: Reporting in Vivo Experiments) guidelines (Kilkenny *et al*., 2010).

Maintenance of adult zebrafish was conducted according to previously described husbandry standard protocols at 28°C with a 14:10 hours (h) light:dark cycle (Lawrence, 2007). The following zebrafish lines were used; *Tg(kdrl:HRAS-mCherry)^s916^* labelling EC membrane (Hogan *et al*., 2009), *Tg(klf2a:GFP)* expressing GFP under control of the zebrafish *klf2a* promoter, *Tg(sm22ab:nls-mcherry)^sh480^* labelling mural cells expressing a smooth muscle actin binding protein and *Tg (nbt:GCaMP3)* which allows quantification of neuronal calcium levels (Meza Santoscoy, 2014; Bergmann *et al*., 2018).

### Glucose, mannitol and SNP treatment

Glucose, mannitol and SNP (Sigma) were prepared in E3 medium to final concentrations of 20mM (glucose and mannitol) or 0.1mM SNP (Chhabria *et al*., 2018). Zebrafish larvae for *in vivo* imaging or immunostaining were incubated in E3 medium containing glucose/mannitol from 4-9 days post fertilization (dpf) and SNP from 8-9 dpf (Chhabria *et al*., 2018).

### Assessment of NO reactivity in the cerebral vessels

Larvae exposed to glucose/mannitol with/without SNP as above were incubated with 2.5 µM Diaminofluorescein-FM (DAF-FM) in DMSO (1%) at 9 dpf for 3 hours at 28°C in the incubator in the dark. Larvae were washed with glucose/mannitol solution to remove excess DAF-FM, and then imaged on the lightsheet microscope.

### Glutamine synthetase, GFAP and TRPV4 Immunohistochemistry

Immunohistochemistry (IHC) was used to observe and quantify changes in TRPV4 (AB2241068; ThermoFisher) and the glial specific markers, GS (mab302; Millipore) and GFAP (zrf-1; Zebrafish International Resource Center, Oregon). The protocol was adapted from (Inoue and Wittbrodt 2011). Larvae for each different treatment (mannitol or glucose, with or without co-treatment with SNP) were fixed in 4% Paraformaldehyde (PFA) overnight at 4°C followed by a 5min wash with 1x PBS before resuspending in 100% Methanol (MeOH) for storage at −20°C. Samples were rehydrated from MeOH with 3x 10min washes with phosphate buffer saline + 0.1% tween (PBST), with gentle agitation on a rocker. Larvae were then suspended in 150mM Tris-HCl (pH 9) for 5 minutes, followed by heating at 70°C for 15 minutes. Larvae then underwent 2x 10min washes with PBST, followed by 5min washes with distilled water (dH2O). Larvae were permeabilized using ice cold acetone at −20°C for 20 minutes, followed by 2x 5min washes with dH2O, then equilibrated 2x 5min washes in PBST. Subsequently, larvae were incubated in blocking buffer (B-buffer: 10% sheep serum, 0.8% triton X100 and 1% bovine serum albumin in PBST) for 3 hours at 4°C. B-buffer was replaced with incubation buffer (I-buffer: 1% sheep serum, 0.8% triton X100 and 1% bovine serum albumin in PBST) containing the primary (1°) antibodies (Abs): 1:250 GS (mouse monoclonal), 1:100 GFAP (mouse monoclonal), 1:300 TRPV4 (rabbit polyclonal) and 1:1000 DAPI followed by incubation at 4°C for 3 days with gentle agitation on a rocker.

Residual 1° Abs were removed by 3x hourly washes in PBST (at room temperature (RT)). Larvae then underwent 2x 10 min washes with PBS + 1% triton X100, followed by 2x hourly equilibration washes in PBS-TS (PBS + 1% triton X100+10% sheep serum).

Larvae were incubated in I-buffer containing secondary antibodies (2° Abs): anti-rabbit, 488 nm (A28181, Invitrogen), anti-mouse, 647 nm (A-11011, Invitrogen) and anti-mouse, 561 nm (A-11034, Invitrogen), corresponding to respective 1° Abs, at 1:500 dilutions for 2.5 days, in the dark on a rocker at 4°C.

Prior to imaging on the lightsheet microscope, larvae were washed three times in PBS-TS (at RT), followed by two 1 hour washes with PBST, each at RT. Larvae were mounted in 1% low melting point agarose (LMP) (Sigma) and imaged for the glial patterning in the brain for different markers.

### Lightsheet fluorescent imaging

Lightsheet fluorescent microscopy (LSFM) was performed on 9 dpf larvae on a Zeiss Lightsheet Z.1 microscope. Larvae were minimally anesthetised (using 4.2% *v/v* tricaine methanesulfonate) and embedded in 1% agarose in a glass capillary (inner diameter ∼ 1mm) while imaging. We acquired 3D z stacks with 800 x 600 pixels (1 pixel = 0.6 µm) in X-Y direction and a depth of 100 slices in Z directions (slice thickness = 1 µm).

For imaging spontaneous neuronal activity, we acquired time lapses of single ‘z’ plane optic tectum with our previous acquisition settings and frequency quantifications (Chhabria *et al*., 2018).

### Image analysis: *klf2a* quantification

Acquired 3D image stacks were converted to 2D maximum intensity projections and the tectal vasculature was segmented out. Tectal vascular length was extracted (Chhabria *et al*., 2018). The segmented vasculature was used as a binary mask, followed by normalizing the total intensity of the green channel (*klf2a:GFP*) in the optic tectum to the tectal vascular length.

### Image analysis: quantification of vascular mural cells

Acquired 3D image stacks were converted to 2D maximum intensity projections followed by segmenting the *sm22ab^sh480^* nuclei (red channel) using intensity based thresholding similar to the method of RBC segmentation described in (Chhabria *et al*., 2018). Segmented cells in the optic tectum were enumerated (in a fixed vascular volume) using custom written MATLAB scripts used for all treatment groups.

### Behavioural analysis

#### Light/Dark preference

Analysis of light/dark preference of the larval zebrafish was designed on a similar principle to that of the adult zebrafish light/dark preference test (Blaser and Penalosa 2011). A 12-well plate was modified by adhering three cellophane films (blue, green and yellow) to half of each well to create a ‘dark’ side that allowed the camera to track larvae movement by infrared (IR). Larvae from different treatment groups were placed on the light side of the well filled with 5 ml E3 (without methylene blue).

The plate was placed inside a Viewpoint-Zebrabox system (1% light intensity) and the tracking protocol was built allocating dark and light regions of each well prior to the start of imaging (to get *x,y* coordinates of dark/light regions separately for analysis). Speed thresholds were set as high speed > 6.4mm/s, low speed 3.3-6.3mm/s, and inactive < 3.3mm/s. Total experimental duration was one hour, inclusive of acclimatization (recording started immediately post adding larvae to individual well).

#### Automated locomotion analysis

Using previous rodent based methods developed for Morris water maze (Wolfer & Lipp, 2000; Graziano *et al*., 2003; Gehring *et al*., 2015; Illouz *et al*., 2016; Vouros *et al*., 2018), we quantified four different features from the swimming trajectories of zebrafish. Coordinates of the swimming trajectories were extracted from the Viewpoint-Zebrabox system and were segmented into smaller paths delimited by light/dark, followed by quantification of path features listed in **Figure 1**. Small path segments, with lengths lower than 1^st^ percentile of segments, generated as an artefact of light/dark transitions were removed from further analysis. The path features eccentricity (ε) and mean point distance from ellipsoid (MPDE) quantify the spatial elongation of the locomotion trajectories and are used as measures of exploration in the field while mean point distance from centre (MPDC) of the well, represents thigmotaxis (preference of edge vs. centre of well). Additionally, we quantified the number of transitions between light and dark areas (**Figure 1**).

**Figure 1:**
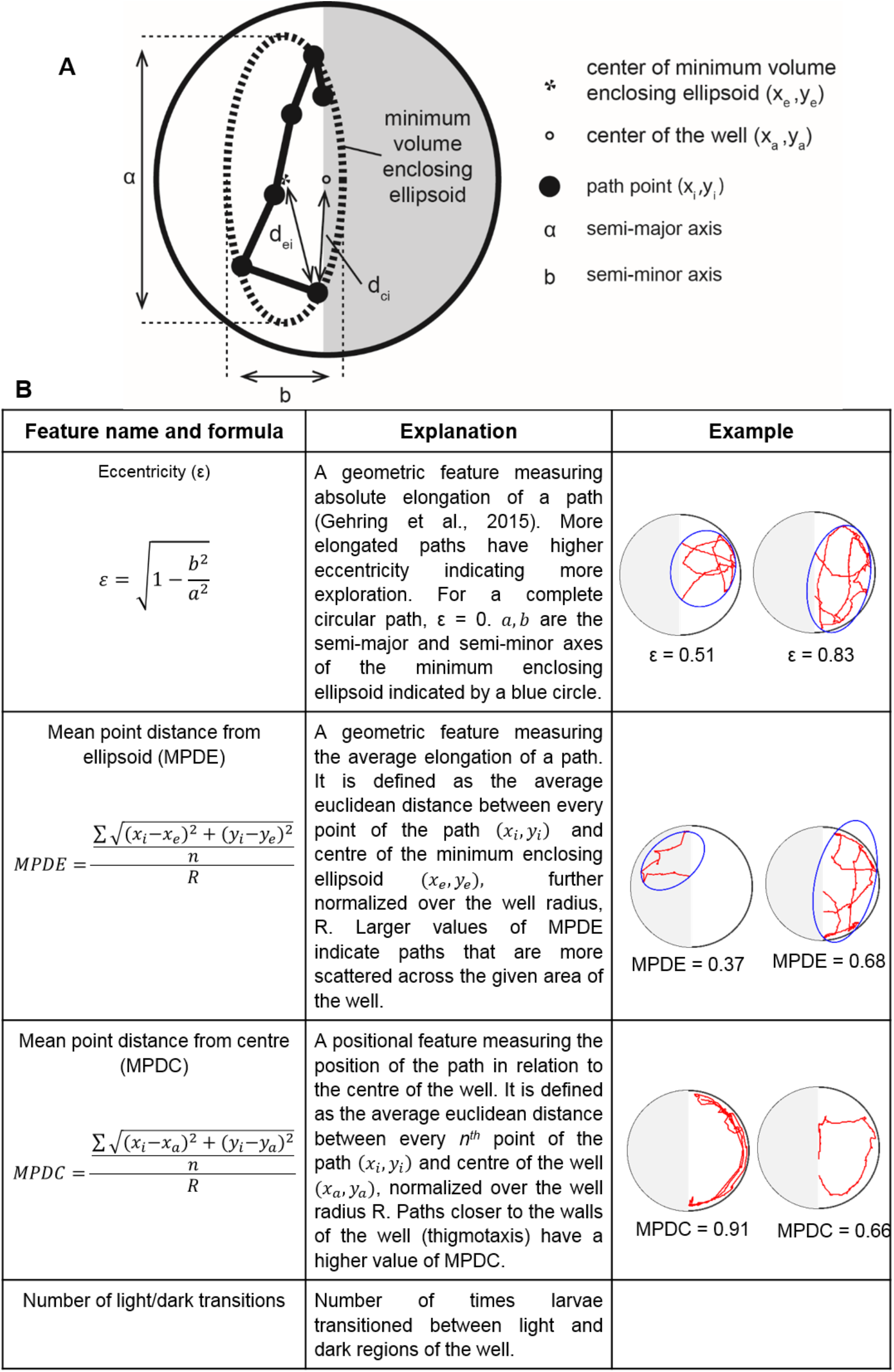
Mathematical description of features calculated from the larval trajectories. **A:** Schematic diagram of the well showing parameters calculated for various features. **B:** Table describing the formulae of calculated features for each coordinates of the trajectories. The minimum enclosing ellipsoid with centre (*x_e_*, *y_e_*) and major and minor axes of *a* and *b* is defined as the unique closed ellipse of smallest volume which enclose all points (*x_i_*, *y_i_*) of a path. 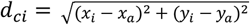 is the Euclidean distance of every point of the path to centre of the well, (*x_a_*, *y_a_*). 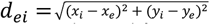 is the distance between the centre of minimum enclosing ellipsoid (*x_e_*, *y_e_*) and *i^th^* point of the path. For each feature, two distinctive numeric examples are provided.

### Experimental design and statistical analysis

Experiments were designed using the NC3Rs experimental design analysis (EDA) tool. GraphPad Prism (La Jolla, CA®) was used for statistical comparisons. All intergroup comparisons were performed using two-way ANOVA with posthoc multiple comparison tests (Sidak’s test) where appropriate. *P* values <0.05 were considered to be statistically significant. Data are shown as mean ± standard error of mean (s.e.m.) unless specified.

## Results

### Glucose exposure reduces vascular nitric oxide reactivity which is prevented by co-treatment with SNP

Studies with diabetic patients have shown reduced bioavailability of NO in the ECs (Williams *et al*., 1996; Pieper, 1998). We therefore first examined whether glucose exposure reduces NO availability in the cerebral vessels. We used DAF-FM staining to visualise NO reactivity in 9dpf zebrafish embryos exposed to 20mM Glucose or mannitol (as osmotic control) with or without co-treatment with 0.1mM SNP. Representative micrographs of the vessels in the left optic tectum are shown in **Figure 2A**. We observed variable levels of NO reactivity that co-localised with the *kdrl:HRAS-mcherry* endothelial reporter in animals treated with mannitol. Co-treatment with mannitol and the NO donor, SNP did not alter the intensity of vascular NO reactivity compared to treatment with mannitol alone (**Figure 2**). Exposure to glucose significantly reduced vascular NO reactivity in the tectal vessels (**Figure 2**), in keeping with data from other models (Pieper, 1998; Du *et al*., 2001). Co-treatment with SNP prevented this reduction in vascular NO reactivity (**Figure 2**).

**Figure 2:**
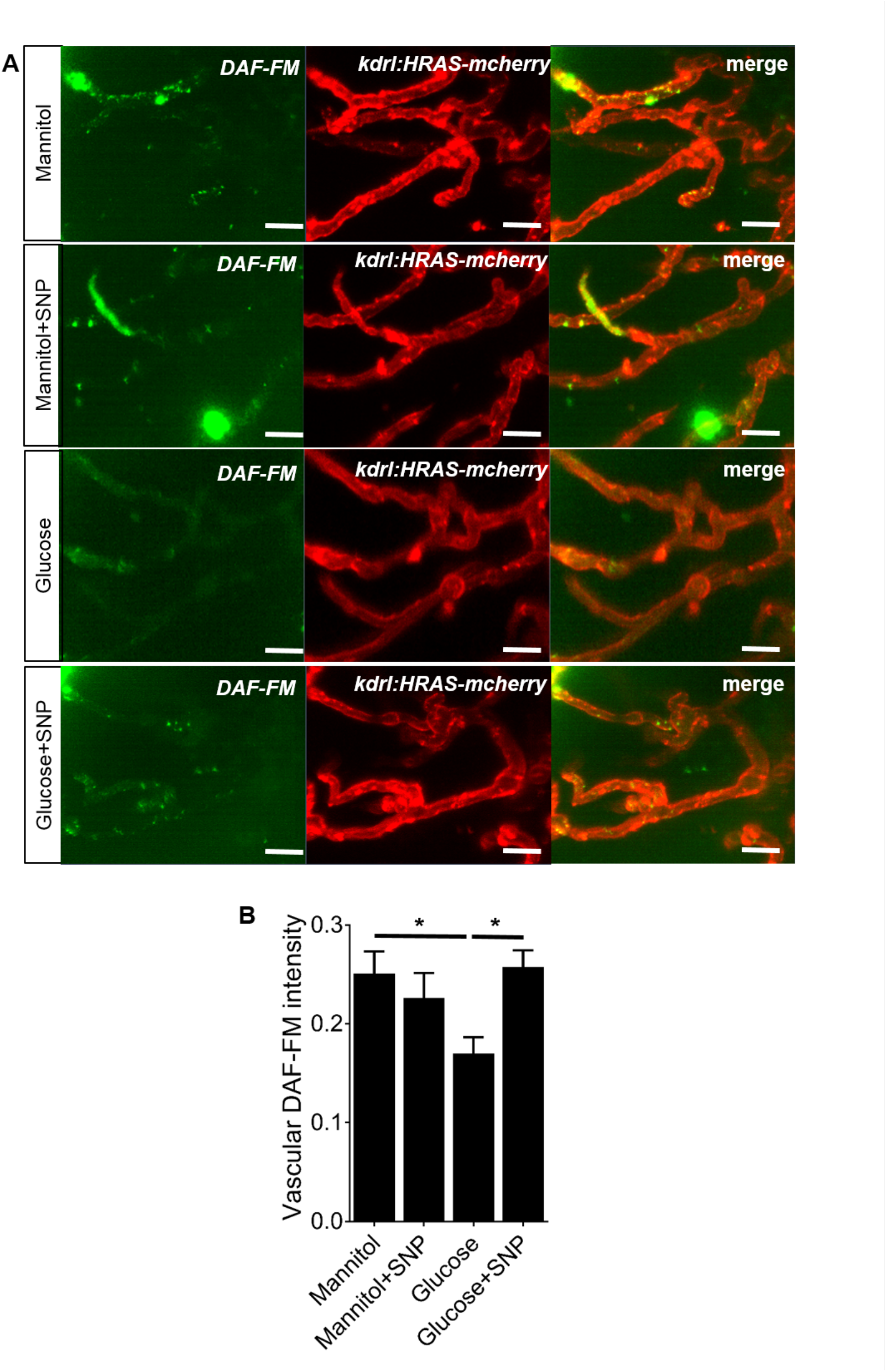
Effect of mannitol/glucose treatment with/without SNP on NO reactivity, quantified by intensity of DAF-FM staining. **A:** Representative micrographs of tectal vessels showing separate and merged channels (green: DAF-FM staining, red: *kdrl:HRAS-mcherry*) for 20mM mannitol or glucose exposed larvae co-treated with or without SNP. Scale bar represents 20 µm. **B:** Quantification of DAF-FM intensity in the tectal vessels (n=25, 24, 27 and 24 larvae for mannitol, mannitol + SNP, glucose and glucose + SNP, respectively). Data are mean ± s.e.m. *p<0.05.

### Glucose exposure reduces endothelial *klf2a* which is prevented by SNP co-treatment

The shear-stress responsive transcription factor *klf2a* is protective against vascular disease (Dekker *et al*., 2002; Dekker *et al*., 2005; Chiu *et al*., 2009). To determine whether glucose exposure affects endothelial shear stress sensing and *klf2a* expression, we quantified the intensity of a *Tg(klf2a:GFP)* reporter in the cerebral vessels of zebrafish with glucose or mannitol with or without SNP co-treatment. Representative micrographs are shown in **Figure 3A**. Glucose exposure significantly reduced intensity of the *klf2a:GFP* reporter compared to mannitol alone (**Figure 3B**). Although SNP co-treatment with mannitol had no effect compared with mannitol alone (**Figure 3**), SNP co-treatment prevented the glucose-induced reduction in the *klf2a* reporter expression.

**Figure 3:**
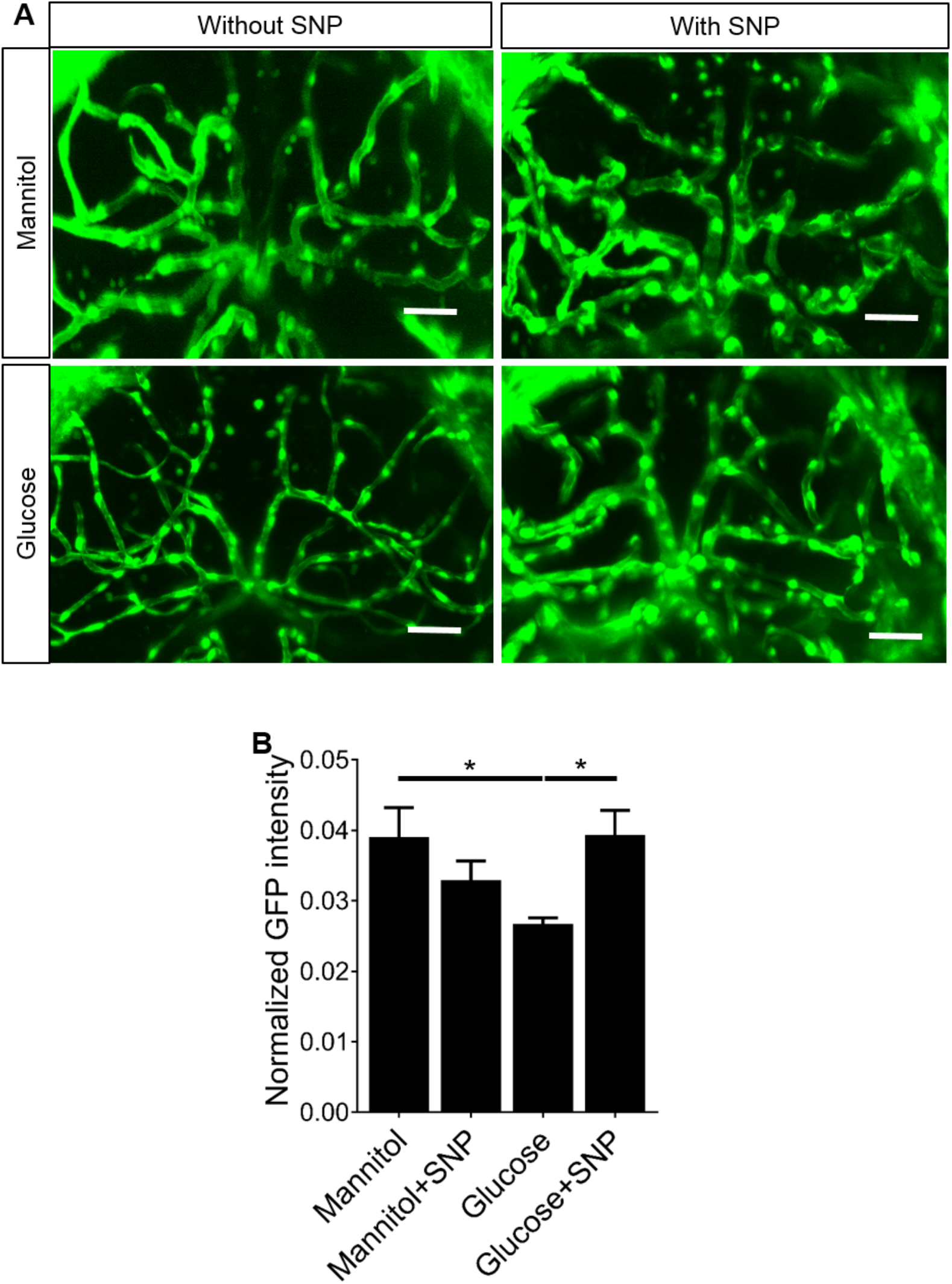
Effect of mannitol/glucose treatment with/without SNP on expression of *klf2a:GFP* expression. **A:** Representative micrographs of tectal vessels in *Tg(klf2a:GFP)* exposed to 20mM mannitol or glucose co-treated with or without SNP. Scale bar represents 20 µm. **B:** Quantification of the *klf2a:GFP* intensity in the tectal vessels (n=26 larvae/group). Data are mean ± s.e.m. *p<0.05.

### Glucose exposure reduces the number of vascular mural cells on the tectal vessels which is prevented by SNP co-treatment

NO is required for mural cell function and can evoke hyperpolarization in mural cells (including pericyte and smooth muscle cell) causing vasodilation (Sakagami *et al*., 2001; Lee *et al*., 2005). Mural cell loss is a feature of diabetes (Pfister *et al*., 2008) but no therapy has been shown to reverse this. We therefore examined the effect of glucose on vascular mural cells. We used a *sm22ab:nls-mCherry* reporter to quantify the number of vascular mural cells present in the optic tectum. Representative micrographs are shown in **Figure 4A**. Glucose exposure induced a significant reduction in the number of vascular mural cells compared with either mannitol or mannitol plus SNP (**Figure 4B**). Co-treatment with SNP prevented the reduction of vascular mural cells induced by glucose exposure.

**Figure 4:**
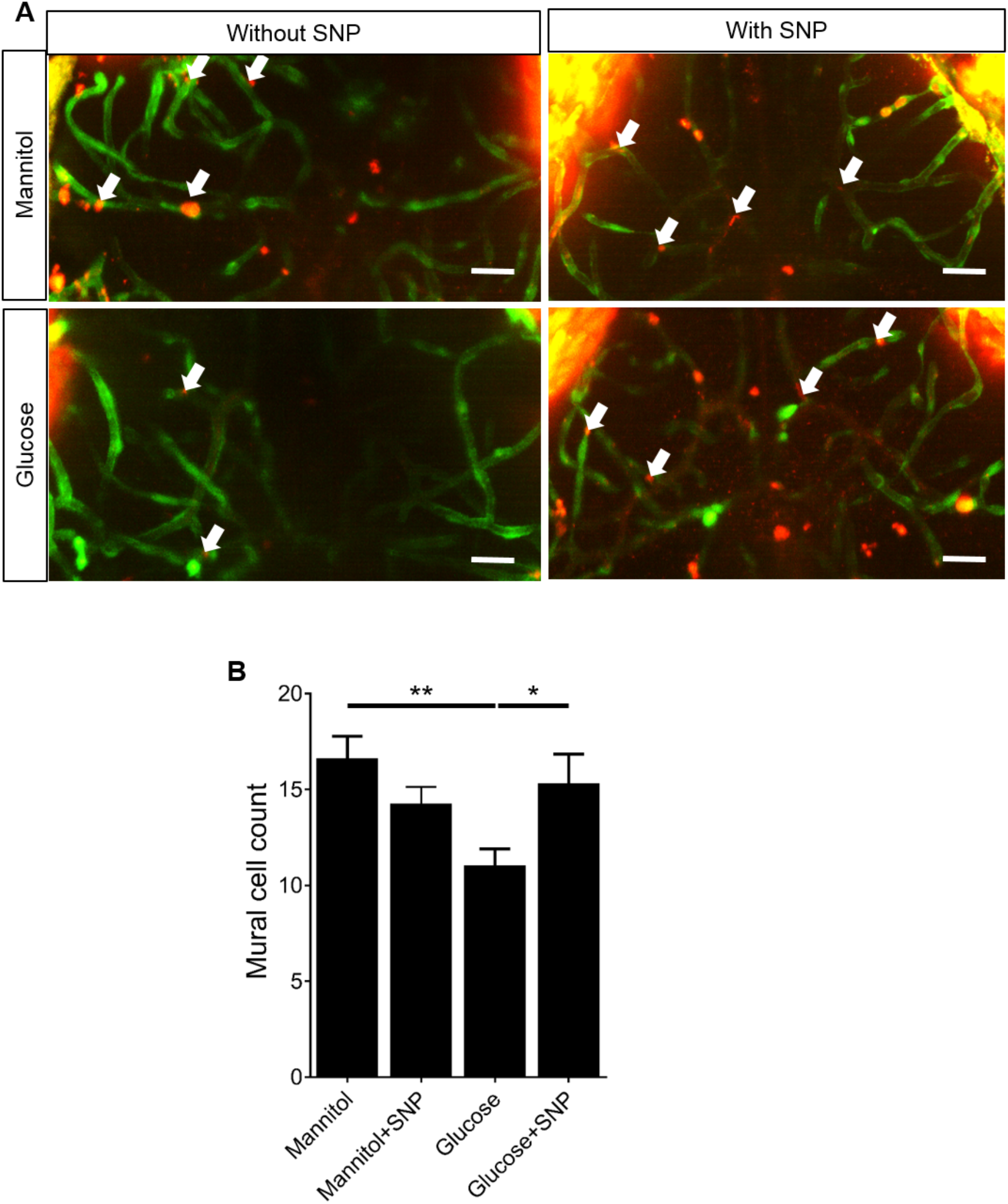
Effect of mannitol/glucose treatment with/without SNP on mural cell number on the tectal vessels. **A:** Representative micrographs of tectal vessels showing separate and merged channels (green: *fli1:GFF:UAS:GCaMP6*, red: *sm222ab:nls-mcherry ^sh480^*) for 20mM mannitol or glucose exposed larvae co-treated with or without SNP. Scale bar represents 20 µm. White arrows indicate mural cell nuclei. **B:** Quantification of the number of *sm22ab:nls-mcherry^sh480^* nuclei on the tectal vessels for 20mM mannitol or glucose exposed larvae co-treated with or without SNP (n=28 larvae/group). Data is mean ± s.e.m. *p<0.05, **p<0.01.

### Glucose exposure increases GFAP expression in the optic tectum which is prevented by SNP co-treatment

Glial cells play major roles in maintenance of the blood brain barrier and NVU function (Janzer & Raff, 1987; Prat *et al*., 2001). Experimental studies have shown over-expression of GFAP (termed astrogliosis) in response to both hyperglycemia and type 1 diabetes (Coleman *et al*., 2004). We thus examined the effect of glucose or mannitol with or without SNP on GFAP expression. Representative micrographs of whole mount 9 dpf old zebrafish are shown in **Figure 5A**. Glucose exposure increased GFAP expression compared to mannitol treatment (**Figure 5B**), in keeping with astrogliosis in other diabetic models. This was prevented by co-treatment with SNP (**Figure 5B**).

**Figure 5:**
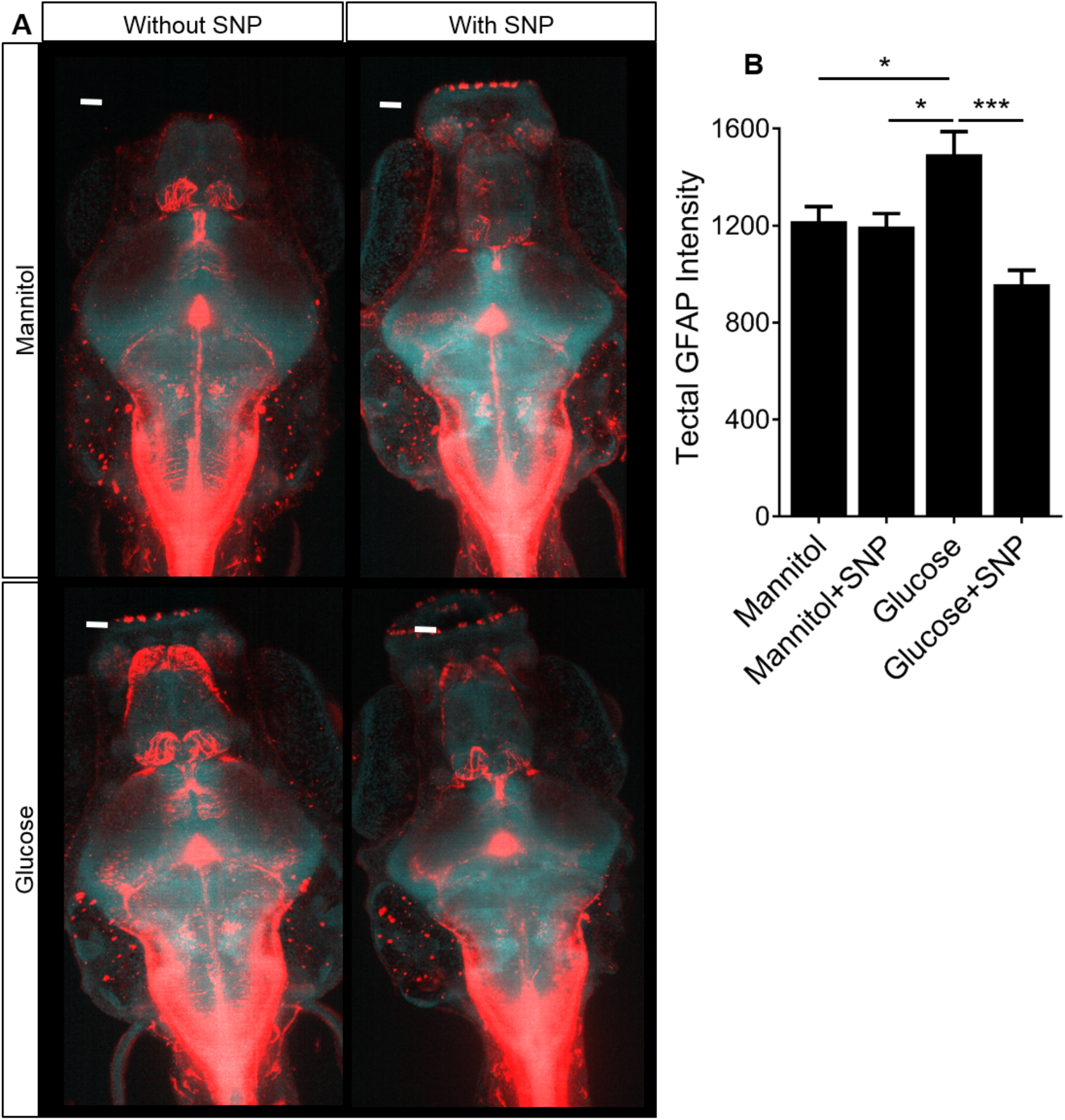
Effect of mannitol/glucose treatment with/without SNP on GFAP expression. **A:** Representative micrographs showing the effect of mannitol/glucose exposure with/without SNP treatment on GFAP expression (Red channel represents GFAP and blue channel represents DAPI). Scale bar represents 20 µm. **B:** Quantification of GFAP expression in the optic tectum (n=16, 12, 18 and 20 larvae for larvae for mannitol, mannitol + SNP, glucose and glucose + SNP, respectively). Data are mean ± s.e.m. *p<0.05, ***p<0.001.

### Glucose exposure reduces the expression of TRPV4 in the optic tectum which is prevented by SNP co-treatment

Since hyperglycemia downregulates TRPV4 in the ECs of the retinal microvasculature (Monaghan *et al*., 2015), we investigated TRPV4 expression by immunohistochemistry in 9 dpf old zebrafish larvae exposed to glucose or mannitol with or without SNP treatment. We performed immunohistochemistry for TRPV4 to first compare expression patterns in the optic tectum. Representative micrographs are shown in **Figure 6**. Glucose exposure decreased tectal TRPV4 expression (which includes radial glial, endothelial or neuronal expression of TRPV4) compared to mannitol. Co-treatment with SNP prevented the effect of glucose on TRPV4 expression (**Figure 6**).

**Figure 6:**
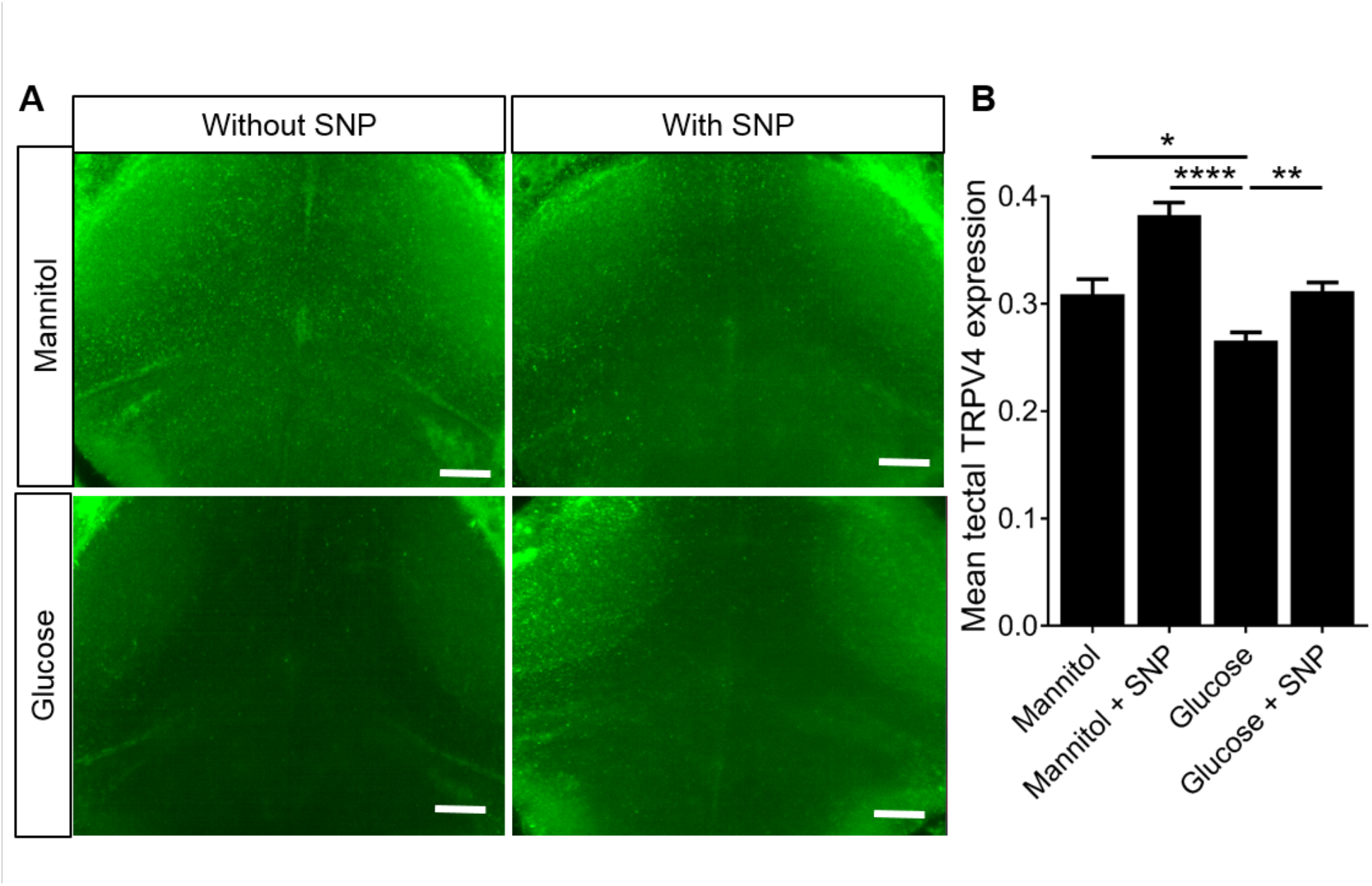
Effect of mannitol/glucose treatment ± SNP on TRPV4. **A:** Representative micrographs of optic tectum showing the effect of mannitol/glucose treatment with/without SNP treatment on the expression of TRPV4. Scale bar represents 20 µm. **B:** Quantification of the TRPV4 intensity in the optic tectum in a fixed volume of the tissue (n=21, 20, 24 and 26 larvae for larvae for mannitol, mannitol + SNP, glucose and glucose + SNP, respectively). Data are mean ± s.e.m. **p<0.01, ****p<0.0001.

### Glucose exposure reduces expression of glutamine synthetase in the optic tectum which is prevented by SNP co-treatment

Glutamine synthetase, a glial-specific enzyme involved in recycling of extracellular/extrasynaptic glutamate to glutamine, is reduced in neurological disorders and diabetes (Lieth *et al*., 2000; Burbaeva *et al*., 2003). We therefore examined expression of glutamine synthetase in the optic tectum. Glucose exposure induced a significant reduction in glutamine synthetase expression, which was prevented by SNP co-treatment (**Figure 7**).

**Figure 7:**
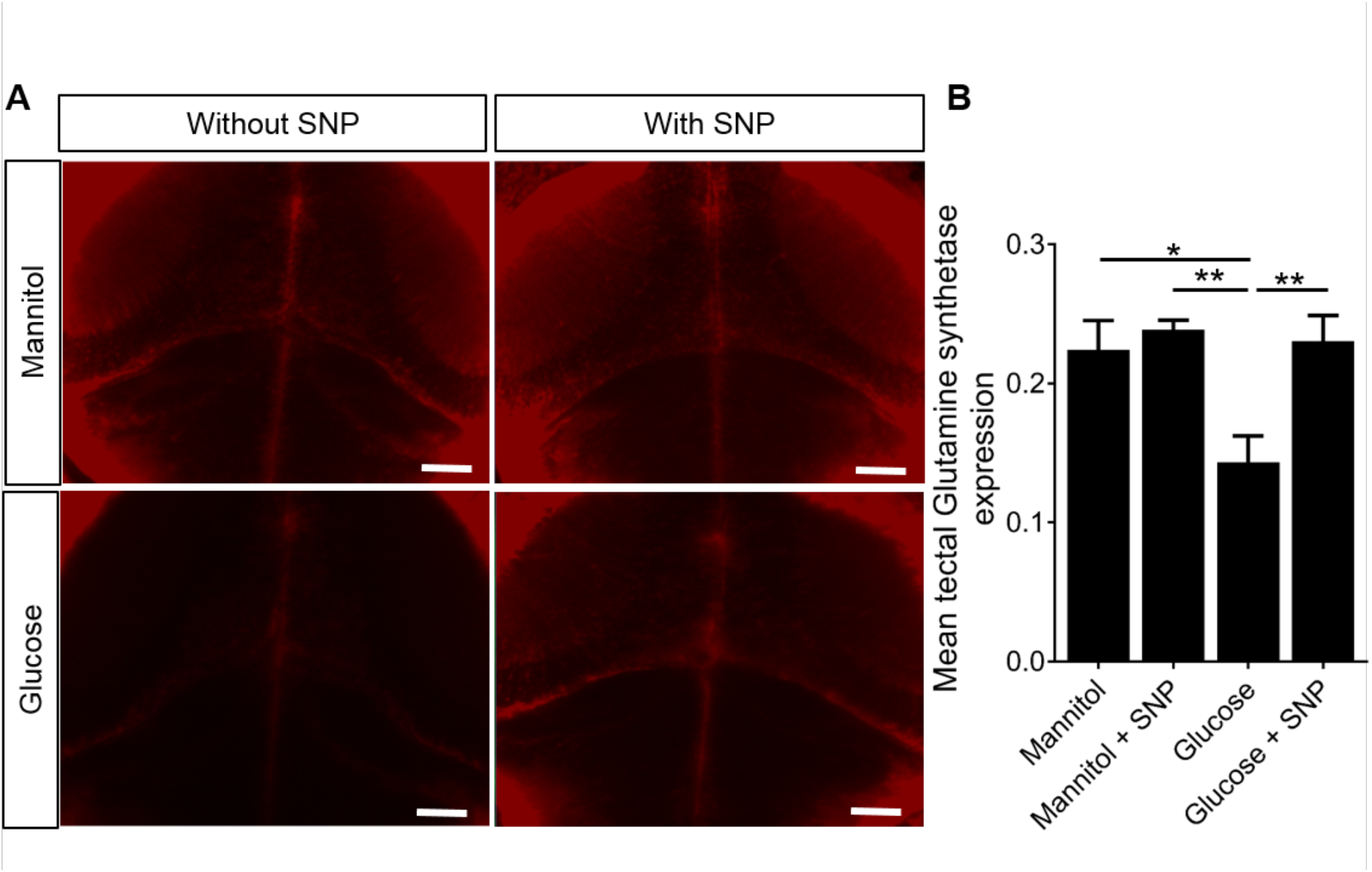
Effect of mannitol/glucose treatment with/without SNP on glutamine synthetase. **A:** Representative micrographs of optic tectum showing the effect of mannitol/glucose treatment with/without SNP treatment on the expression of glutamine synthetase. Scale bar represents 20 µm. **B:** Quantification of the glutamine synthetase intensity in the optic tectum in a fixed volume of tissue (n=21, 20, 24 and 26 larvae for larvae for mannitol, mannitol + SNP, glucose and glucose + SNP, respectively). Data are mean ± s.e.m. *p<0.05, **p<0.01.

### Glucose exposure increases frequency of spontaneous neuronal calcium transients which is prevented by SNP co-treatment

The previous results showed that glucose exposure affects both the anatomy of the NVU (vascular mural cell loss) and induces dysregulation of gene expression, including a reduction of glutamine synthetase which might be expected to cause neuronal hyperexcitability. We therefore quantified neuronal activity in our model. Representative time series of spontaneous neuronal activation (quantified as *ΔF/F*_o_) in larvae exposed to mannitol or glucose with or without SNP are shown in **Figure 8A**. Glucose exposure increased neuronal activation compared to mannitol exposure (**Figure 8B**). Co-treatment with SNP prevented this glucose-induced increase in neuronal activation (**Figure 8B**), suggesting that diabetes-induced neuronal hyperexcitability may be mediated via reduced NO bioavailability.

**Figure 8:**
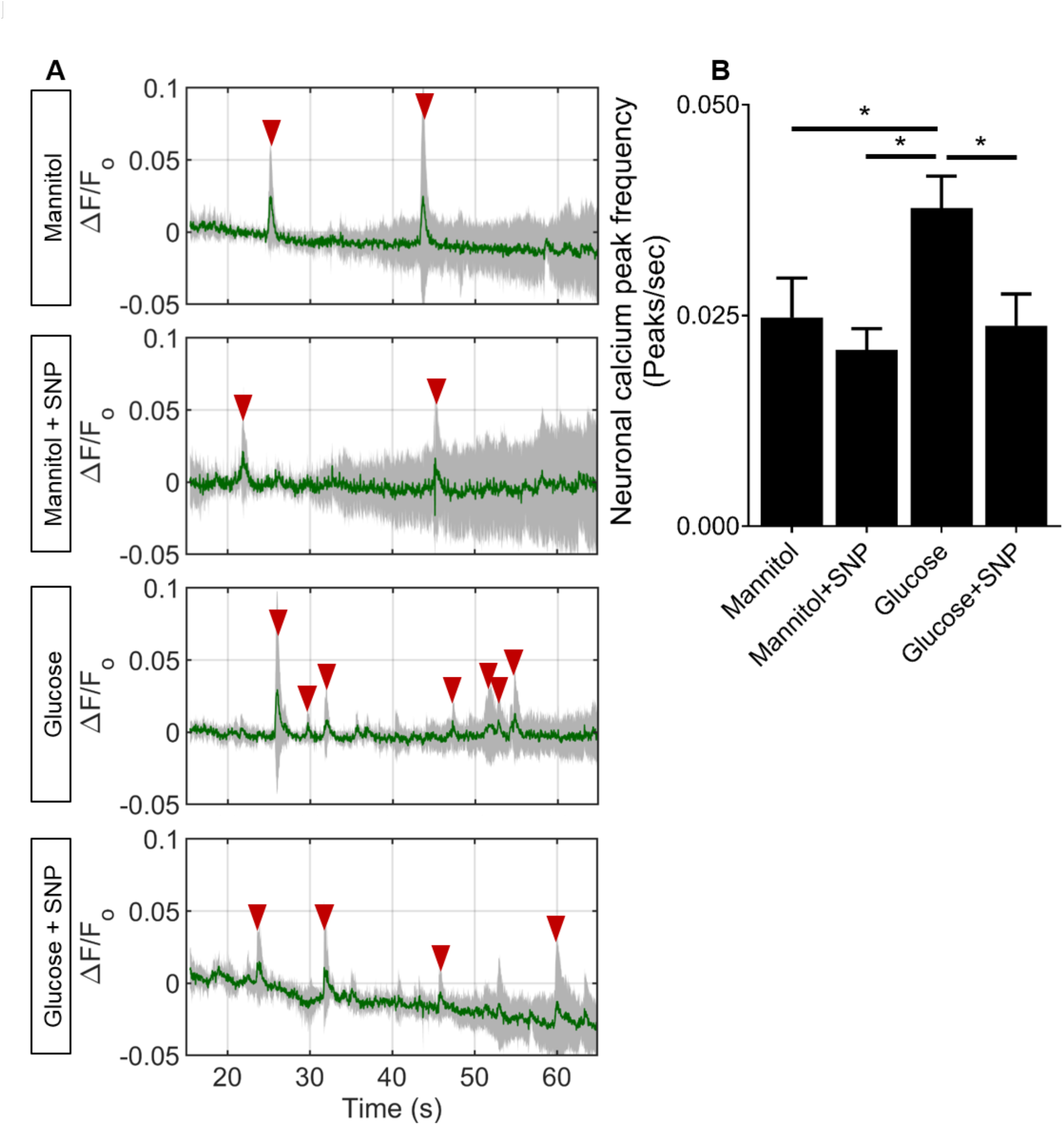
Effect of mannitol/glucose treatment with/without SNP on frequency of spontaneous neuronal calcium transients. **A:** Time series of neuronal activation (*ΔF/F_o_* in zebrafish (5 larvae/group) exposed to (left:top-to-bottom); mannitol, mannitol+SNP, glucose and glucose+SNP. Red arrowheads indicate the detected peaks. **B:** Quantification of frequency of neuronal calcium transients for each of the groups (n=28 larvae/group). Data are mean ± s.d. in **A** and mean ± s.e.m in **B**. *p<0.05

### Glucose exposure alters light/dark preference and locomotion which is prevented by SNP co-treatment

Although the previous data clearly demonstrate molecular and functional defects of the NVU induced by glucose, if such disturbances are clinically relevant, they would be expected to manifest in overt behavioural or neurological consequence. We therefore assessed the effect of glucose exposure on free swimming behaviour in zebrafish larvae to attempt to identify behavioural consequences of glucose exposure. We first examined the effect of glucose exposure on light-dark preference. **Figure 9A** shows the trajectories of representative zebrafish larva for a period of 1 h in each treatment group. Red and green paths indicate high and low speed locomotion respectively. We tested light/dark preference using percentage of time spent in the light or dark side of the well. Mannitol exposed larvae at 9 dpf showed a preference for light, spending ∼80% time in the light. This was significantly reduced by glucose exposure (**Figure 9B**). However, co-treatment with SNP prevented the effect of glucose on light/dark preference (**Figure 9B**).

**Figure 9:**
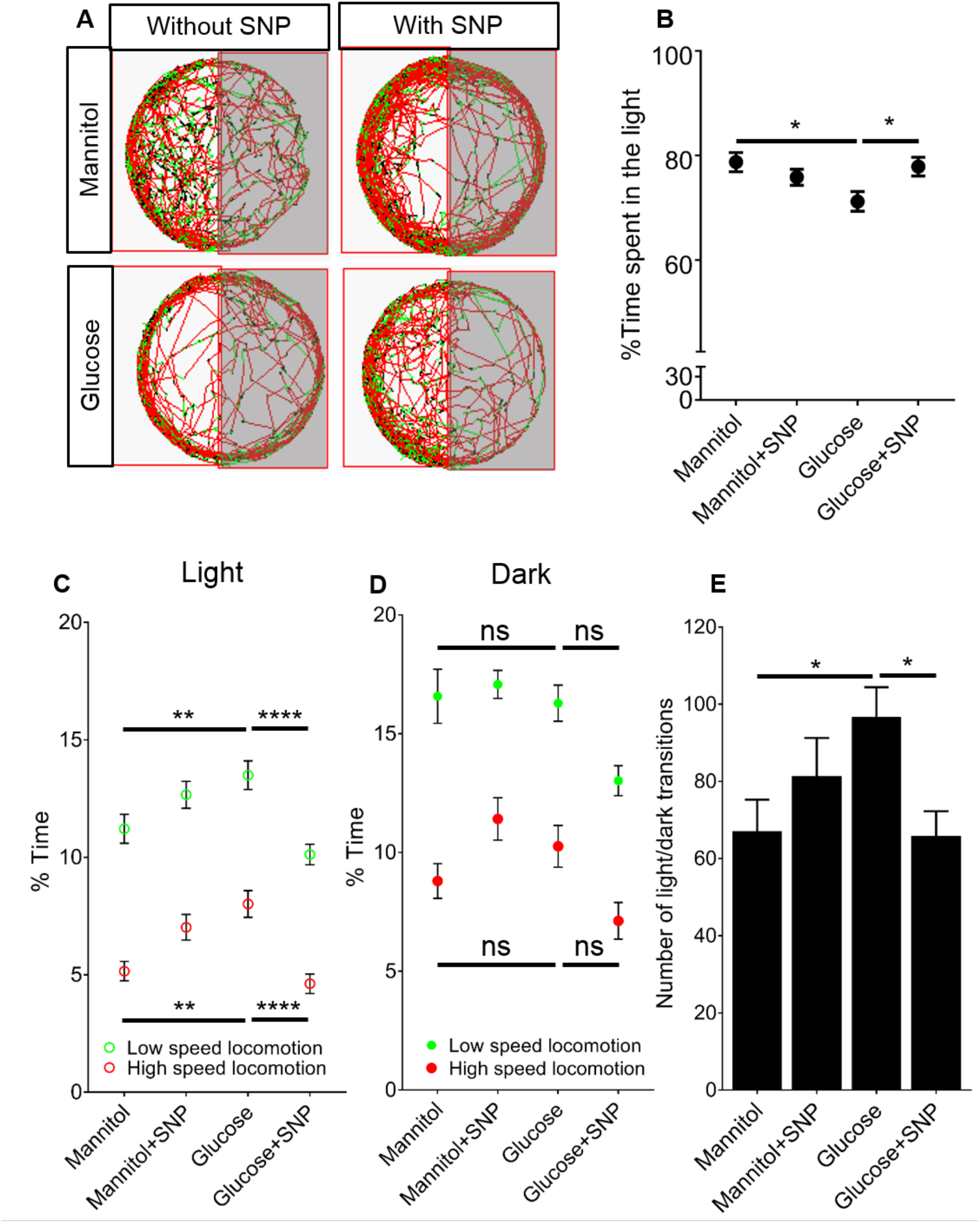
Effect of mannitol/glucose treatment with/without SNP on larval zebrafish behaviour. **A:** Representative trajectories of 9 dpf old zebrafish moving in half darkened wells (of a 12 well plate) as tracked by Viewpoint software for mannitol or glucose with or without SNP treatment. Red trajectories represent high speed (> 6.4mm/s), green low speed (3.3-6.3 mm/s), and black inactive (< 3.3mm/s) **B:** Percentage of time spent in light region of the well by larvae (n = 50, 45, 44 and 56 larvae for mannitol, mannitol+SNP, glucose and glucose+SNP, respectively). **C:** Quantification of percentage time spent in low and high speed locomotion in the light region by the same animals as in B. **D:** Quantification of percentage time spent in low and high speed locomotion in the dark region by the same animals as **B-C. E:** Quantification of number of transitions into the light/dark regions for same larvae as in B-D. Data are mean ± s.e.m. *p<0.05, **p<0.01 and ****p<0.0001.

To further characterise larval behaviour, we quantified the time spent in high speed (> 6.4mm/s), low speed (3.3-6.3 mm/s), and inactive < 3.3mm/s locomotion in the light (**Figure 9C**) or dark side of the wells (**Figure 9D**). Glucose exposure significantly increased the time larvae spent in both low and high speed locomotion while in the light and SNP prevented this effect (**Figure 9C**). In contrast, glucose did not induce any significant difference in the amount of time spent in either high or low speed locomotion while in the dark side of the well (**Figure 9D**). We quantified the number of transitions between light and dark areas of the well and found that glucose exposure induced a significant increase in the number of transitions that was prevented by co-treatment with SNP (**Figure 9E**).

We next investigated three further features of larval zebrafish locomotion; path eccentricity, mean point distance to ellipsoid (MPDE), and mean point distance to centre (MPDC) (**Figure 1**) in the light and dark sides of the wells. Glucose induced significant increases in path eccentricities, MPDE and MPDC in the light side of the well which was prevented by SNP (**Figure 10A-C**). When we quantified these behaviours in the dark side of the well, we saw a similar significant increase in path eccentricity and MPDE but not in MPDC (**Figure 10D-F**). Again, SNP prevented the effect of glucose on these aspects of behaviour.

**Figure 10:**
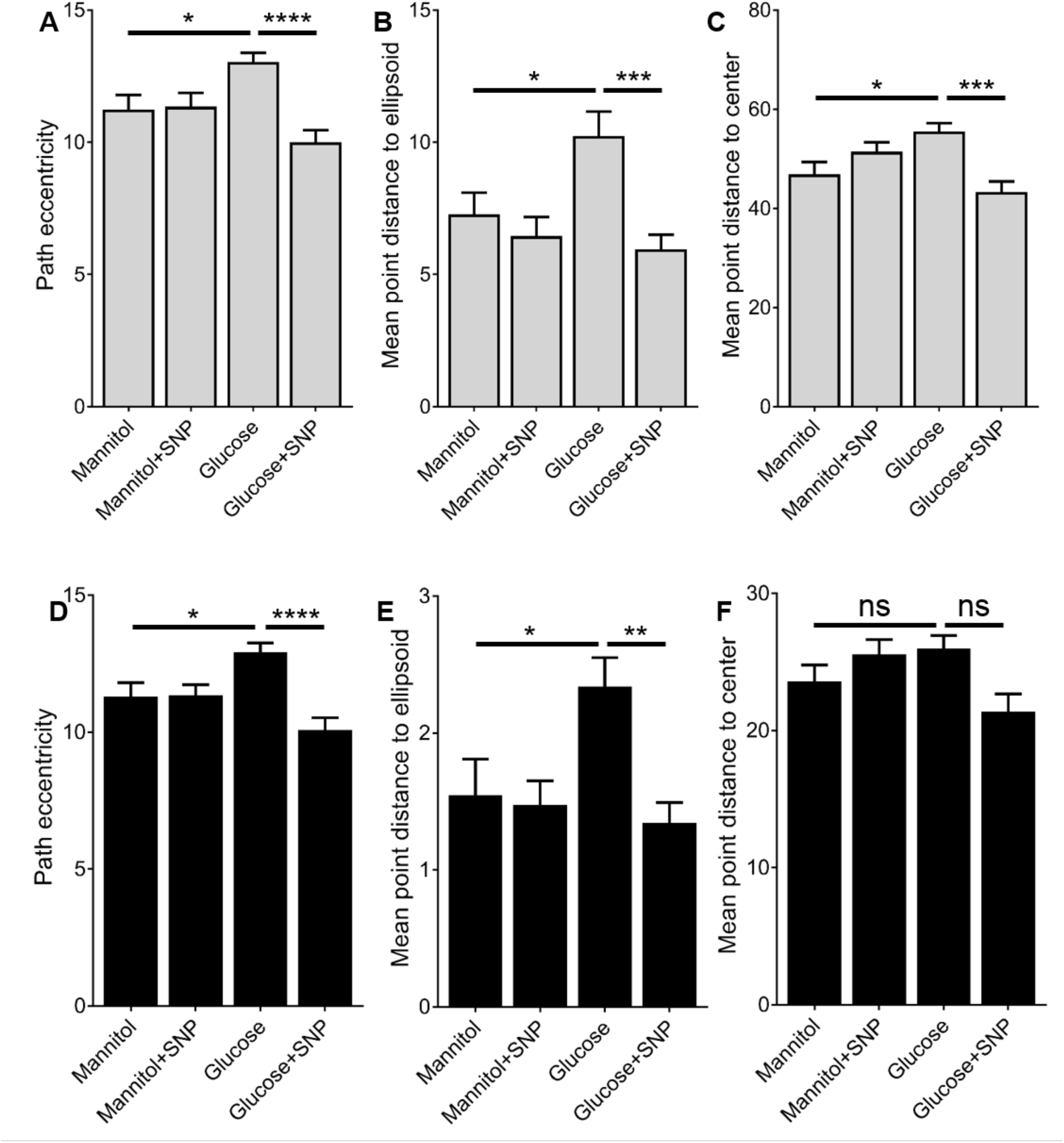
Effect of mannitol/glucose treatment with/without SNP on various features of zebrafish locomotion. Quantification of mean frequency of path eccentricities **(A, D)**, MPDE **(B, E)** and MPDC **(C, F)** (n= 50, 45, 44 and 56 larvae for mannitol, mannitol+SNP, glucose and glucose+SNP, respectively) in **A-C:** light and **D-F:** dark regions of the well. Data are mean ± s.e.m. *p<0.05, **p<0.01, ***p<0.001, ****p<0.0001

## Discussion

We here describe a comprehensive molecular, anatomic, functional, and behavioural study of the effect of glucose exposure on the NVU in zebrafish. The zebrafish model has a range of advantages, coupling simplicity, speed, and cost-effectiveness with sophisticated *in vivo* imaging in a whole-organism setting. We show that a relatively short (5d) exposure to 20mM glucose (a concentration seen in the blood of poorly controlled human diabetics (Amiel *et al*., 1988; Boyle *et al*., 1988)) affects every constituent cell type of the NVU that we examined. Glucose exposure impaired both vascular NO production and vascular mural cell number, which builds on our previous work showing that tectal endothelial patterning is impaired by glucose exposure (Chhabria *et al*., 2018). These effects were also associated with reduced endothelial *klf2a* expression, which is known to promote vascular inflammatory gene expression, thrombosis, and atherosclerosis (SenBanerjee *et al*., 2004a; Lin *et al*., 2005). Developmental studies in *klf2a(-/-)* mice have shown reduced vascular mural cell recruitment suggesting the role of *klf2a* in maintaining endothelial-mural cell interactions (Wu *et al*., 2008; Fukuhara *et al*., 2009; Gaengel *et al*., 2009). Abnormal mural cell recruitment or migration is associated with various microangiopathies and is commonly observed in diabetes (Hammes *et al*., 2002; Gaengel *et al*., 2009).

In addition to the negative effects of glucose on the vascular component of the NVU, we found that glucose exposure induced upregulation of GFAP, indicating astrogliosis, with concomitant reductions in glutamine synthetase and TRPV4. In both rodents and zebrafish, TRPV4 channels are expressed on astrocytes, neurons and ECs (Vriens *et al*., 2005; Benfenati *et al*., 2007; Grant *et al*., 2007; Mangos *et al*., 2007; Marrelli *et al*., 2007; Amato *et al*., 2012). Interestingly studies have shown that TRPV4 in ECs can regulate eNOS (Sukumaran *et al*., 2013) and the presence of a feedback loop from eNOS to TRPV4 to modulate TRPV4 based Ca^2+^ signalling (Yin *et al*., 2008). Rodent models have shown that TRPV4 on cortical astrocyte endfeet can evoke changes in intracellular astrocyte calcium concentration, thereby modulating vascular tone and contributing to NVC (Dunn *et al*., 2013; Filosa *et al*., 2013). Rodent experiments support our observations by showing TRPV4 downregulation in streptozotocin induced diabetes (Monaghan *et al*., 2015).

Retinal studies with streptozotocin-induced hyperglycemia in rodents have suggested that hyperplasia of the Muller cells (retinal analog of astrocytes) could lead to an increase in GFAP (Newman & Reichenbach, 1996; Rungger-Brandle *et al*., 2000). Astrocytic glutamate clearance is also impaired under high glucose conditions (Coleman *et al*., 2004) making neurons susceptible to depolarization, a possible cause of neurotoxicity. This could result in accumulation of glutamate in the extrasynaptic space leading to recurrent neuronal depolarization. This is concordant with our observation of an increased number of spontaneous calcium peaks in the glucose treated larvae. Neuronal hyperexcitability and increased firing is known to be associated with seizures, commonly observed in diabetic patients (Martinez & Megias, 2009; Baviera *et al*., 2017). The various cellular markers shown here to be affected by glucose exposure could explain a predisposition of diabetics to seizures. Increased neuronal firing could also lead to abnormal and non – precise pre and post synaptic neuronal firing causing defects in the synaptic plasticity mechanisms necessary for cognition and memory. Further exploration of this could help define the relationship between diabetes and cognitive defects.

The anatomic and molecular effects of glucose exposure on the NVU were associated with altered embryonic behaviour, with a reduction in preference for light, which is a measure of unconditioned anxiety and related disorders in rodents and zebrafish (Kulesskaya & Voikar, 2014; Kysil *et al*., 2017). Unconditioned anxiety is influenced by environmental, emotional and cognitive factors (Arrant *et al*., 2013). It is based on an approach/avoidance conflict between the drive to explore a novel area and an aversion to brightly lit/completely dark open spaces in adult/larval zebrafish, respectively (Bourin & Hascoet, 2003; Arrant *et al*., 2013). The approach/avoidance conflict is well-studied in mammals and is known to have various neural substrates in the brain, including the limbic system, anterior cingulate cortex, ventral striatum and prefrontal cortex (Aupperle *et al*., 2015). Although zebrafish do not possess a cortex, their ventral and dorsal telencephalic area (Vd and Dm, respectively) are homologous to the mammalian amygdala and striatum (Maximino *et al*., 2013). Thus impaired light/dark preference as described in the present study could imply an abnormal circuitry in the zebrafish Vd/Dm. Anatomical studies with zebrafish have shown that Vd/Dm system project to the optic tectum and hence any defects could also affect the optic tectum (Scott & Baier, 2009; Nevin *et al*., 2010).

Various studies have described larval zebrafish behavioural differences with anxiolytic or anxiogenic treatments (Egan *et al*., 2009; Richendrfer *et al*., 2012). However, this is the first study characterizing the effect of hyperglycemia on geometrical and positional characteristics of zebrafish locomotion. Altered positional and geometric features with glucose exposure implies an altered behaviour such as increased exploration and thigmotaxis. Various zebrafish studies have shown increased exploration and thigmotaxis with anxiogenic drug treatments (Egan *et al*., 2009; Blaser *et al*., 2010). This further points to the association of glucose exposure and diabetes to anxiety related brain activation, which warrants further investigation. Although human diabetes is a complex disorder and our zebrafish model examines only the effect of hyperglycemia, our findings broadly reproduce those in other cell based and mammalian model (Williams *et al*., 1996; Li *et al*., 2002). Diabetes is well known to reduce vascular NO levels, and our work reproduces this. Although zebrafish are not known to possess endothelial nitric oxide synthase (eNOS) (Syeda *et al*., 2013), our results strongly indicate vascular NO production. Previous studies have linked hyperglycemia and pharmacologically induced diabetes to reductions in cerebral blood flow (Dandona *et al*., 1978; Stevens *et al*., 2000). A recent study demonstrated rescuing effects of the NO donor, SNP, on hyperglycemia induced neurovascular uncoupling (Chhabria *et al*., 2018). Using the same protocol as described in (Chhabria *et al*., 2018) to induce hyperglycemia in larval zebrafish, we have now described multiple effects of hyperglycemia on cellular markers of the NVU, essential for regulation of CBF and on zebrafish behaviour.

This is the first study to demonstrate that SNP reverses the negative consequences of hyperglycemia on neurovascular anatomy and behaviour. The ability of SNP to prevent all the observed anatomical, molecular and behavioural effects of glucose exposure is exciting as it may represent a possible treatment for diabetes-associated neurovascular dysfunction. Future studies are needed to assess whether these effects of glucose exposure and SNP or other NO donors are seen in mammalian models or humans. NO donors are already widely used clinically for angina and heart failure, and are very cheap and off-patent. Therefore, if mammalian pre-clinical studies support our findings clinical studies could rapidly be performed to examine the ability of NO donors to ameliorate or prevent the consequences of diabetes on neurological dysfunction.

## Author contributions

C.H. and T.J.A.C conceived the work detailed, prepared the manuscript and wrote the grant funding the work. K.C. performed all experiments, performed all data analysis and prepared the manuscript. K.P. assisted with experiments. A.V. and E.V. performed and assisted with behavioural data analysis. C.G. generated the *klf2a* reporter line. Z.J. and R.N.W. generated the *Tg(sm22ab:nls-mcherry)^sh480^* reporter line. R.B.M. provided the antibodies, protocols and assisted with the immunohistochemistry. All authors assisted with writing and editing the manuscript.

## Acknowledgments

We are very grateful to the aquarium staff of the Bateson Centre for expert husbandry and advice. We are grateful to John P. Ashton and Sarah Baxendale for expert training and assistance with the behavioural experiments.

## Funding Acknowledgements

This work was funded by a Project Grant from the National Centre for Replacement, Refinement, and Reduction of Animals in Research (NC3Rs) NC/P001173/1. C.H. is the recipient of a Sir Henry Dale Fellowship jointly funded by the Wellcome Trust and the Royal Society (Grant Number 105586/Z/14/Z). The Zeiss Z1 lightsheet microscope was funded via British Heart Foundation Infrastructure Award IG/15/1/31328 awarded to T.C.

